# Kv1 channels regulate variations in spike patterning and temporal reliability in the avian cochlear nucleus angularis

**DOI:** 10.1101/2021.10.08.463546

**Authors:** James F. Baldassano, Katrina M. MacLeod

## Abstract

Diverse physiological phenotypes in a neuronal population can broaden the range of computational capabilities within a brain region. The avian cochlear nucleus angularis (NA) contains a heterogeneous population of neurons whose variation in intrinsic properties results in electrophysiological phenotypes with a range of sensitivities to temporally modulated input. The low-threshold potassium conductance (G_KLT_) is a key feature of neurons involved in fine temporal structure coding for sound localization but a role for these channels in intensity or spectrotemporal coding has not been established. To determine whether G_KLT_ affects the phenotypical variation and temporal properties of NA neurons, we applied dendrotoxin (DTX), a potent antagonist of Kv1-type potassium channels, to chick brain stem slices in vitro during whole-cell patch clamp recordings. We found a cell-type specific subset of NA neurons were sensitive to DTX: single-spiking NA neurons were most profoundly affected, as well as a subset of tonic firing neurons. Both tonic I (phasic onset bursting) and tonic II (delayed firing) neurons showed DTX sensitivity in their firing rate and phenotypical firing pattern. Tonic III neurons were unaffected. Spike time reliability and fluctuation sensitivity measured in DTX-sensitive NA neurons was also reduced with DTX. Finally, DTX reduced spike threshold adaptation in these neurons, suggesting that G_KLT_ contributes to the temporal properties that allow coding of rapid changes in the inputs to NA neurons. These results suggest that variation in Kv1 channel expression may be a key factor in functional diversity in the avian cochlear nucleus.

**New and noteworthy:** The dendrotoxin-sensitive voltage-gated potassium conductance typically associated with neuronal coincidence detection the timing pathway for sound localization is demonstrated to affect spiking patterns and temporal input sensitivity in the intensity pathway in the avian auditory brain stem. The Kv1-family channels appear to be present in a subset of cochlear nucleus angularis neurons, regulate spike threshold dynamics underlying high-pass membrane filtering, and contribute to intrinsic firing diversity.

## Introduction

Neurons in early sensory nuclei must process a wealth of incoming information to extract features that are relevant to their particular circuit. The avian cochlear nucleus is composed of two divisions, nucleus magnocellularis (NM) and nucleus angularis (NA), each of which extract unique information from auditory nerve activity in parallel (1). NM neurons encode timing information by phase-locking to the fine structure in the acoustic signal (2, 3), partly enabled by specialized intrinsic properties such as a large low threshold voltage-gated potassium conductance (G_KLT_) (4–8). This dendrotoxin (DTX)-sensitive conductance is attributed to the expression of the Kv1 family of potassium channels, particularly Kv1.1 and Kv1.2, that are prominent throughout the auditory brainstem of birds and mammals (9–15).

In contrast to NM, NA is classically understood to process information about sound intensity or level in service of interaural level difference computation. (3, 16–18) More broadly, however, NA neurons are also involved in spectrotemporal and amplitude modulation coding and auditory envelope processing.(19, 20) The neuronal population in NA shows greater morphological and physiological heterogeneity than those in NM and NL (1, 21, 22). Sound-evoked responses observed in vivo in NA fall into a range of response categories, such as onset, chopper, primary-like, and “Type IV”, similar to corresponding recorded responses in the mammalian cochlear nucleus (2, 23, 24). Likewise, NA neurons display a range of electrophysiological phenotypes that likely arise from the expression of a more diverse array of ion channels (22, 25–29). Single spiking neurons in NA most closely resemble NM neurons, while a broad group of repetitively firing neurons in NA can be further subdivided according to their firing patterns to step currents in vitro: early burst firing (tonic I), late or delayed firing (tonic II) and regular spiking (tonic III) (25, 26). A DTX-sensitive conductance has been shown to be required for the single-spiking phenotype, but blocking this conductance also elevated firing rates in tonic neurons (22). Along with immunohistochemical evidence for Kv1 channel protein expression in NA (10, 30), these data suggest that G_KLT_ is present and active in shaping NA neuron physiology. The ion channel basis of the different tonic phenotypes is not yet known.

The parallels between the heterogeneity among in vivo response types and the heterogeneity among in vitro response types suggests that intrinsic properties may have a significant effect on the transformation of auditory information. We have recently investigated type-specific variation in sensitivity to temporally modulated stimuli in the tonic neurons in vitro and shown how diversity in intrinsic properties can enhance stimulus encoding (26, 27). Temporal modulation sensitivity in vitro and bandpass feature extraction in vivo could be replicated in NA neurons using a phenomenological model of adaptive spike thresholds (19, 28). While the model was originally derived from the biophysical phenomenon of sodium channel inactivation (31), its ion channel basis in NA has not been empirically confirmed and could also be influenced by other subthreshold conductances, including G_KLT_ (32–34).

In this study, we used whole cell patch clamp electrophysiology and DTX to pharmacologically block G_KLT_ and test whether G_KLT_ was needed to elicit the various spiking phenotypes among the tonic neurons. We further investigated whether G_KLT_ was necessary for the temporal reliability of firing, high-pass filtering properties, and spike threshold adaptation that characterized a subset of tonic firing neurons.

## Methods & Materials

### Brain slice preparation

All animal procedures were performed with Institutional Animal Care and Use Committee approval and according to University of Maryland guidelines on animal welfare. Chicken (*Gallus gallus*) embryos incubated to embryonic day 17-18 were cooled and rapidly decapitated, and the head section containing the brain stem blocked, placed in a chilled, oxygenated low-sodium artificial cerebrospinal fluid (ACSF)(low-Na^+^ ACSF in mM: 97.5 NaCl, 3 KCl, 2.5 MgCl_2_, 26 NaHCO_3_, 2 CaCl_2_, 1.25 NaH_2_PO_4_, 10 dextrose, 3 HEPES, and 230 sucrose) and dissected out of the cranium. The brain stem tissue block was mounted with cyanoacrylate glue and supported with 5% agarose gel solution. Transverse slices (250 μm thick) containing NA were cut on a vibrating tissue slicer (Leica Microsystems, Wetzler, Germany). Slices were incubated in normal ACSF (in mM: 130 NaCl, 3 KCl, 2 MgCl_2_, 26 NaHCO_3_, 2 CaCl_2_, 1.25 NaH_2_PO_4_, 10 dextrose, and 3 HEPES) at 34°C for 30 min then held at room temperature in normal ACSF until recording.

### Whole cell patch-clamp electrophysiology

Slices were placed in a recording chamber and continuously perfused with oxygenated, warmed normal ACSF (1–2 mL/min, 28-30°C) containing synaptic blockers (3 ***μ***M strychnine, 20 ***μ***M SR95531 (gabazine), 15 ***μ***M DXQX, and 20 ***μ***M AP5; Sigma) to isolate intrinsic activity. Whole cell patch-clamp recordings were performed on visually identified NA cells using infrared differential interference contrast video microscopy. Initial glass recording pipette resistances were 3–7 MΩ. Pipettes were filled with a potassium gluconate intracellular recording solution (in mM: 110 potassium gluconate, 20 KCl, 1 EGTA, 2 MgCl_2_, 10 HEPES, 2 Na_2_ATP, 0.3 Na_2_GTP, 10 phosphocreatine and 0.2% biocytin). Series resistance, cell capacitance, and resting membrane voltage were measured upon break-in in voltage clamp mode. Voltage recordings were made with a MultiClamp 700B amplifier (Molecular Devices, Sunnyvale, CA) set to current-clamp mode. Application of the current stimulus and recording of the voltage output were controlled by an analog-to-digital board (National Instruments, Austin, TX) and a computer running custom software written in IGOR Pro (WaveMetrics, Lake Oswego, OR). A holding current was applied to maintain a constant voltage baseline of approximately -60 mV. Drug application for pharmacological manipulation of voltage response was performed by bath application (100 nM **α**-dendrotoxin, Tocris-Cookson).

Cell type classification protocols were carried out to identify neuronal phenotypes in NA with 400-ms-duration flat current steps of varying amplitudes from −150 pA up to 750 pA in 50 or 100 pA intervals. We collected data from 77 NA neurons (20 single-spike, 12 tonic I, 16 tonic II, 23 tonic III, and 6 damped). The patterns of action potential (AP) firing over multiple current steps were used to divide NA neurons into 3 broad groups: single-spiking, tonic firing or damped. Single spiking NA neurons resembled neurons in NM and NL, firing a single onset AP at all step amplitudes. Tonic firing neurons fired repetitively with overshooting action potentials throughout the duration of the flat current steps at most amplitudes, but could be further subdivided based on their characteristic firing at depolarizing steps just above rheobase: burst firing at the onset of current injection (tonic I), a delay followed by burst firing (tonic II), or tonic firing with uniform interspike intervals (tonic III) (see (25) for more details). Damped neurons fired broad APs that progressively declined in amplitude into oscillatory subthreshold potentials. If the neuron type was ambiguous from the current levels utilized, subsequent, smaller intervals were used to distinguish the phenotype. Passive membrane properties were assessed using a small hyperpolarizing current step (−50 pA).

### Acquisition of f-I curve and analysis

Input-output functions were assessed by measuring firing-current (f-I) curves with standard flat current steps (400 ms duration as for classification protocols) or noisy current stimuli (2 second duration). Stimuli to acquire noise *f*-I curves were constructed by convolving Gaussian white noise with an exponential function (time constant, 3 ms) added to a DC step function as described previously (26, 27). These noise stimuli simulate the arrival of many small, stochastic, and statistically independent synaptic currents, both excitatory and inhibitory. To accommodate cell-to-cell differences in input resistance, a standard noise current was created by calibrating the standard deviation (around a zero mean) of the noise current to generate a 2-mV standard deviation in the membrane potential in the target neuron, designated “1σ”. The amount of voltage fluctuation was varied by multiplying the standard noise stimulus by a factor of 2, 4, or 8 (i.e. 4-, 8-and 16-mV voltage fluctuation, respectively). Each trial had an interstimulus interval of 8-10 seconds. A hyperpolarizing conditioning pre-pulse (−50 pA, 1 sec) was applied prior to each stimulus to minimize sodium channel inactivation from the prior stimulus. A complete series of stimuli in the parameter space [mean DC, noise level] was systematically generated by varying the mean current amplitude and noise level independently (3 noise levels, 5-7 DC levels). Firing rates were averaged across 3 stimulus repetitions over the full 2-second duration and reported as mean ± SD in Hertz. After acquisition of control data, DTX (100 nM) was bath-applied to the slice and data for drug trials acquired after a 10 minute wash in period.

Metrics to quantify differences between the lowest noise level (2σ) and highest noise level (8σ) curves were devised. One metric quantified the total area difference index (ADI)(28):

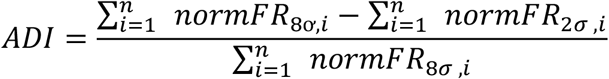

where normFR was the normalized firing rate at each of *n* steps in each noise level. The second metric was the normalized difference in the maximum firing rate between 2σ and 8σ, ΔMaxFR:

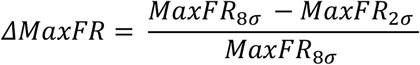

where MaxFR refers to the maximum firing rate value of each *f*-I curve. Both metrics varied from 0 (curves are identical) to 1 (maximum difference). Plotting these points (MaxFR index and the ADI index) against each other produced a spectrum of fluctuation sensitivity. Neurons closest to the origin were more “integrator-like”, while neurons further away from the origin were more “differentiator-like”. Using the criteria of ADI and ΔMaxFR >0.2, we were able to test the noise *f*-I curves from 15 differentiators and 5 integrators with DTX. ΔMaxFR index and ADI index were statistically compared using Wilcoxon’s t-test.

### Spike timing reliability

After noise was calibrated, a 4σ noisy current was injected into neurons at a single DC current step amplitude (typically 150-400 pA) to produce a firing rate between 20-50 Hz. Repeated trials of 2 second long frozen noisy current injections were injected (usually 30 to 45 trials, or until >600 spikes were collected) before and after DTX application. To quantify neuronal reliability, we calculated a shuffled autocorrelogram (SAC), a histogram of all interspike intervals between spikes across trials, excluding intervals within the same trial. The SAC is normalized by the normalizing factor (NF):

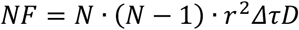

where N is the number of trials, r is the average firing rate, Δτ is the bin width of the correlation function (0.2 ms) and D is the length of the stimulus (2 sec) (26, 27, 35, 36). The SAC was fit with a Gaussian function, whose central peak is defined as the correlation index (CI). The precision of the firing between trials is represented in the full-width at half-maximum (FWHM) of the Gaussian curve, measured by

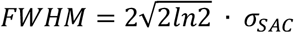

where *σ*_*SAC*_ represents 1 standard deviation of the gaussian fit of the SAC. Spike threshold was determined using the first derivative of the voltage signal and applied uniformly for all trials. CI and FWHM, referred to as width, were quantified and compared using Wilcoxon’s t-test.

### Mutual information

To quantify the mutual information (I) between spike trains under control conditions (X) versus those recorded in DTX (Y), spike times were binned (1 ms) and their time bin distributions (x_i_, y_i_) were used to calculate Shannon’s equation for mutual information (37, 38) :

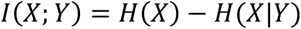

where H(X) is the marginal entropy, or the observable information of X:

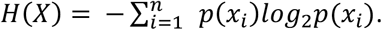

where *p(x*_*i*_) is the probability of distribution x_i_ and the entropy is given in bits. H(X|Y),the conditional entropy, is the reduction of uncertainty about X given Y:

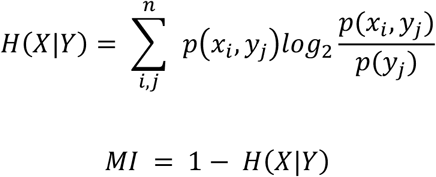

Where p(x_i_,y_i_) is the probability that X = x_i_ and Y = y_i_. To control for random changes or response drift over the recording time, the experimental mutual information was compared to the mutual entropy calculated between control and “sham” conditions, where ACSF without DTX was washed into the recording chamber. Mutual information is reported in bits but bounded by zero and 1, and then subtracted from 1 which allows for zero to mean X and Y are completely independent and 1 to mean X is entirely predicted by Y, analogous to a similarity measure.

### Action Potential Threshold Analysis

The action potential threshold was defined as the voltage at which an all-or-none regenerative spike was initiated. This threshold was determined by using the first derivative of the voltage as previously described (19). A criterion value of the derivative was selected and applied for the entire set of voltage responses across the *f–I* curve. Neither small changes in the criterion value nor the alternative of using the second derivative affected the results of within-subject analyses. Direct comparisons of absolute threshold between neurons, however, were omitted, as differences in the criterion applied could affect across-subject comparisons. The mean, median, and variance of the spike threshold measured across the action potentials for a given stimulus were calculated for each DC step amplitude m and noise level s combination. The mean and median were nearly identical in all cases, as expected for nearly normal distributions.

## Results

### Spiking patterns in NA are shaped by a DTX-sensitive potassium conductance

To determine whether low threshold potassium channel activation was involved in shaping the electrophysiological phenotypes observed in the chick cochlear nucleus NA, we recorded the voltage responses before and after bath application of DTX (100 nM), a specific antagonist of the Kv1.1 and Kv1.2 channels. We characterized the firing patterns and passive membrane properties using a series of flat depolarizing current injections during whole-cell patch-clamp recordings from NA neurons. As shown previously, in control conditions NA neurons displayed heterogeneous spiking phenotypes in vitro that could be broadly classified as single spiking (Fig. 1Ai) or tonic firing (Fig. 1Bi-Di)(25–27) Using small current steps, tonic firing neurons can be further distinguished into 3 subgroups based on whether they tend to show early bursts (tonic I, Fig. 1Bi), late bursts (tonic II, Fig. 1Ci), or repetitive firing evenly spaced for the duration of the stimulus (tonic III, Fig. 1Di)(see also 25).

**Fig. 1.**
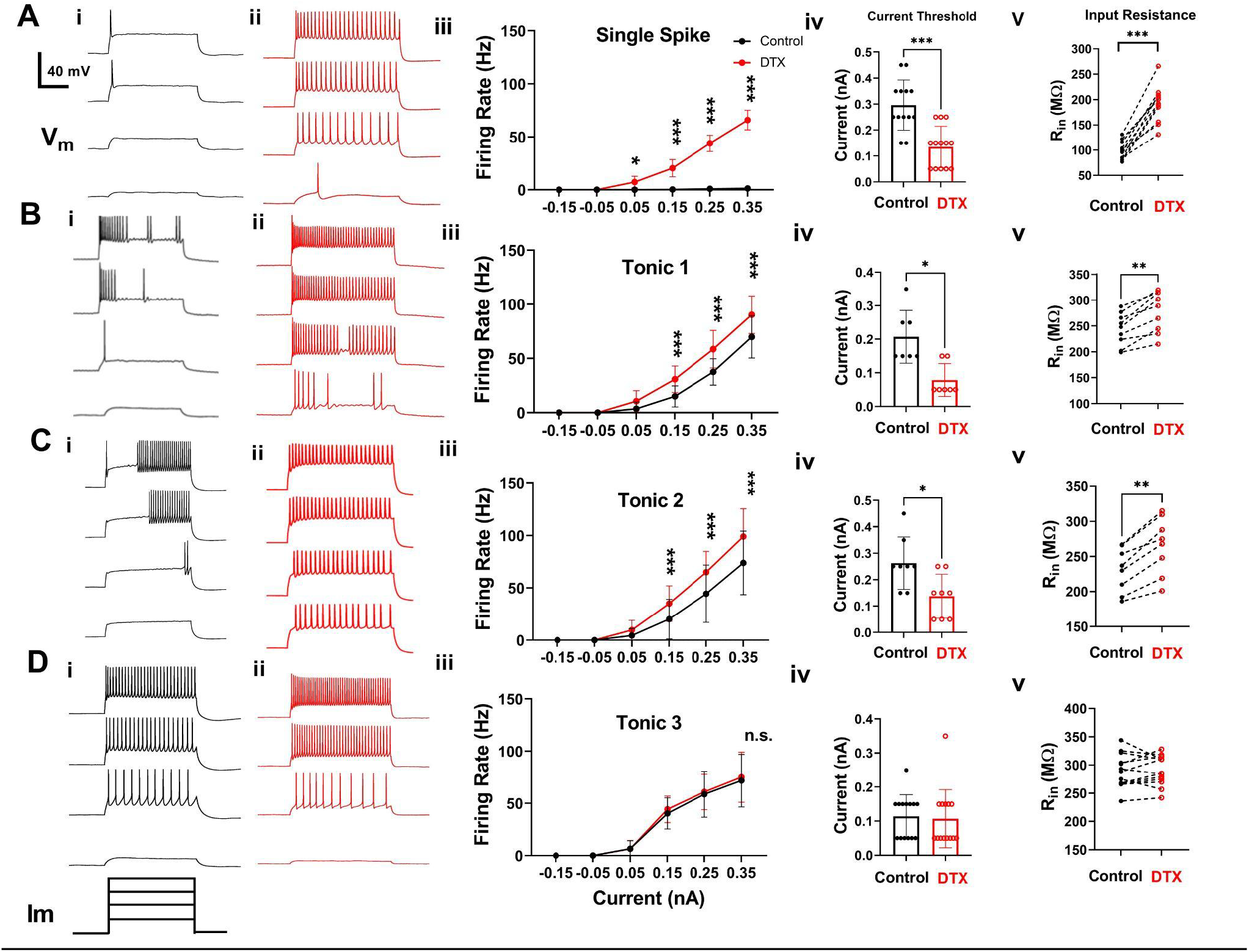
Dendrotoxin altered the spiking responses in a subset of NA neurons. Voltage responses were elicited by a set of current injections of increasing amplitude into NA neurons comprising 4 major cell types: single-spiking (row A) and 3 subtypes of tonic firing neurons: tonic I (row B), tonic II (row C), and tonic III (row D). Example traces show matched responses during control conditions (Ai-Di) and following DTX application (Aii-Dii). In DTX, firing rate increased in single-spike, tonic I, and tonic II neurons (Aiii: P < 0.001; Biii: P < 0.001, Ciii: P < 0.001, 2-way ANOVA) but not in tonic III neurons (Diii, P = 0.25). Asterisks indicate Sidak’s post hoc multiple comparisons test significance, * P < 0.05, *** P < 0.001). The current amplitude threshold (rheobase) was also significantly reduced (Ai: P < 0.001; Biv: P = 0.03; Civ: P = 0.03, Wilcoxon’s signed rank test) except in tonic III neurons (Div, P > 0.99). Input resistance increased in single-spike, tonic I, and tonic II neurons (Av: P = 0.001; Bv: P = 0.039, and Cv: P =0.042, Student’s t-test) except in tonic III neurons (Dv: P > 0.99).

When DTX was applied, changes in the firing patterns were observed in single spiking, tonic I and tonic II neurons. In all single spiking neurons, which characteristically fire single onset spikes upon depolarization in control conditions, DTX application induced repetitive firing throughout the current injection (n = 12; Fig 1Aii), in agreement with a previous study (22). In tonic I neurons DTX application eliminated the burst cessation in these neurons that occurred at low-amplitude steps in control and instead induced firing throughout the step duration at all step levels, (n = 9, Fig 1Bii). Similarly, in tonic II neurons DTX application eliminated the delay in firing at low-amplitude steps in control (n = 8, Fig. 1Cii). Tonic 3 neurons, in contrast, showed no apparent changes in firing pattern (n = 13, Fig. 1Dii).

Changes in the firing patterns also resulted in significant changes in the overall input-output functions for tonic I and tonic II neurons, as well as single spiking neurons (single spiking: n = 12, p = 0.04 at step amplitude 0.05 nA, P < 0.001 at 0.15-0.35 nA, Fig. 1Aiii; tonic 1: n = 9, P < 0.001 at 0.15-0.35nA, Fig. 1Biii; tonic 2:, n = 8, P < 0.001 at 0.15-0.35nA, Fig1Ciii; Sidak’s multiple comparisons test). These firing rate changes were accompanied by a reduction in the threshold current amplitude (rheobase) (Fig. 1A*iv*-C*iv* single spiking control mean = 0.29 ± 0.09 nA, DTX mean = 0.13 ± 0.08 nA, P=0.001, tonic I control mean = 0.21 ± 0.07 nA, DTX mean = 0.78 ± 0.4 nA, P=0.03, tonic II control mean = 0.26 ± 0.07 nA, DTX mean = 0.14 ± 0.8 nA, P = 0.03, Wilcoxon’s test). These changes in excitability were likely partly due to the increase in input resistance measured in these neurons (single-spike: n = 12, control R_in_ = 101.6 ± 17.14 MΩ (mean ± SD), DTX Rin = 191.7 ± 31.37 MΩ, P = 0.001, Fig. 1Av; tonic I: n = 9, control R_in_ = 243.7 ± 29.69 MΩ, DTX R_in_ = 277.8 ± 36.93, P = 0.039, Fig. 1Bv; tonic II: n = 8, control R_in_= 230.6 ± 29.72 MΩ, DTX R_in_ = 265.5 ± 38.1MΩ, P = 0.042, Fig. 1Cv). In contrast, tonic III neurons showed no effects of DTX application on their firing rate, input output function or rheobase (Fig. 1D*i*-*iv* tonic III control mean rheobase = 0.11 ± 0.06 nA, DTX mean rheobase= 0.10 ± 0.08 nA, P > 0.99) nor in their input resistance (n = 13, Fig. 1Dv, control R_in_ = 289.7 ± 26.2 MΩ, DTX R_in_ = 291.0 ± 24.9 MΩ, P = 0.47, Fig. 1Div). Finally, a separate subset of NA neurons were characterized in control conditions as having a ‘damping’ phenotype: broader action potentials that diminished in amplitude and whose voltage oscillates around a membrane voltage. In these neurons, DTX application had no apparent effect on the voltage response (Fig. 2A). Neither the damping constant (Fig. 2B) nor the input resistance (Fig. 2C) were altered. Together, these results indicate that a DTX-sensitive conductance, the putative G_LKT_ conductance, contributes to the spiking patterns and firing rates of a specific subset of NA neurons.

**Fig. 2.**
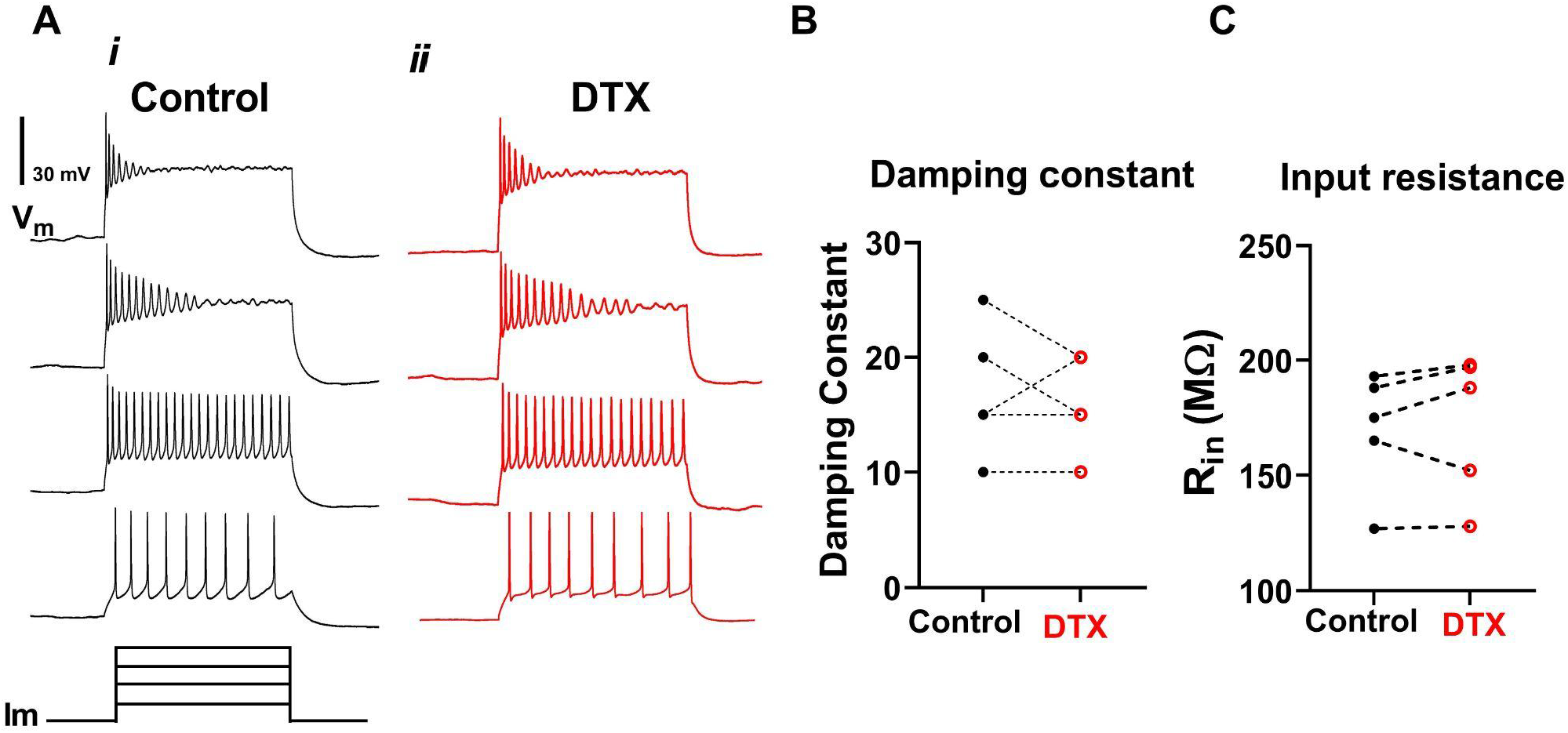
DTX did not alter firing or intrinsic properties of damped neurons. Voltage responses in response to flat current injections (400 ms duration) showed no apparent change between control (Ai) and after DTX application (Aii). The damping constant (B) and input resistance (C) were unchanged (control damping constant = 19.0 ± 6.3 (mean ± SD), DTX damping constant = 18.2 ± 5.2, P = 0.97; control R_in_ = 168.3 ± 22.4 MΩ, DTX R_in_ = 174.8 ± 26,7MΩ, P > 0.99, Student’s t test).

### The DTX-sensitive conductance improves temporal firing reliability and information content in tonic firing neurons

Different subtypes of NA neurons exhibited varying levels of temporal reliability in response to frozen noise stimulation in vitro (26). To test whether G_KLT_ regulated temporal reliability in NA, we collected voltage responses to repeated trials of frozen noise from 49 NA neurons before and after DTX application (9 tonic I, 8 tonic II, 16 tonic III, 5 damped, 11 single-spike) (Fig. 3A, C). The trial-to-trial temporal firing reliability was analyzed using a shuffled autocorrelogram (SAC), a histogram of all across-trial interspike intervals excluding within-trial spike pairs (Fig. 3B) (26, 35). The peak (correlation index, CI) and width of the SAC indicate the degree and precision of the time locking of the spikes across trials. Application of DTX significantly reduced the SAC peak amplitudes measured in single spiking, tonic I and tonic II neurons (single spiking: control CI = 47.4 ± 12.6, DTX CI = 22.4 ± 12.3, n = 11, P = 0.005; tonic I: control CI = 28.9 ± 6.7, DTX CI = 11.1 ± 5.4, n = 9, P =0.0059; tonic II: control CI = 13.7 ± 4.6, DTX CI = 7.0 ± 2.4, n = 8, P = 0.042)(Fig. 3B, C; summary data in Fig. 4A, B) and increased the SAC width (single spiking: control FWHM = 0.5 ± 0.34 ms, DTX FWHM = 1.2 ± 1.0 ms, P < 0.001; tonic I: control FWHM = 2.02 ± 0.93 ms, DTX FWHM = 3.11 ± 1.22 ms, P < 0.001; tonic II: control FWHM = 1.97 ± 0.32 ms, DTX FWHM = 3.41 ± 0.71 ms, P = 0.039, Fig. 4C). In contrast, in tonic III or damped neurons, DTX application had no significant effect on the SAC peak (tonic III: CI = 4.83 ± 3.82, DTX CI = 4.68 ± 3.98, n = 12, P =0.84; damped: control CI = 3.59 ± 1.04, DTX CI = 4.96 ± 2.88, n = 9, P = 0.84) or width (tonic III: FWHM = 3.85 ± 1.64 ms, DTX FWHM = 4.32 ± 2.13 ms, P = 0.41; damped: control FWHM = 6.43 ± 3.14 ms, DTX FWHM = 6.01 ± 3.55 ms, P = 0.56). These results indicate that G_KLT_ contributes to precise and reliable firing in the same subset of cell types that showed firing rate changes with DTX (single spiking, tonic I and II) but not in the tonic III or damped neurons.

**Fig. 3.**
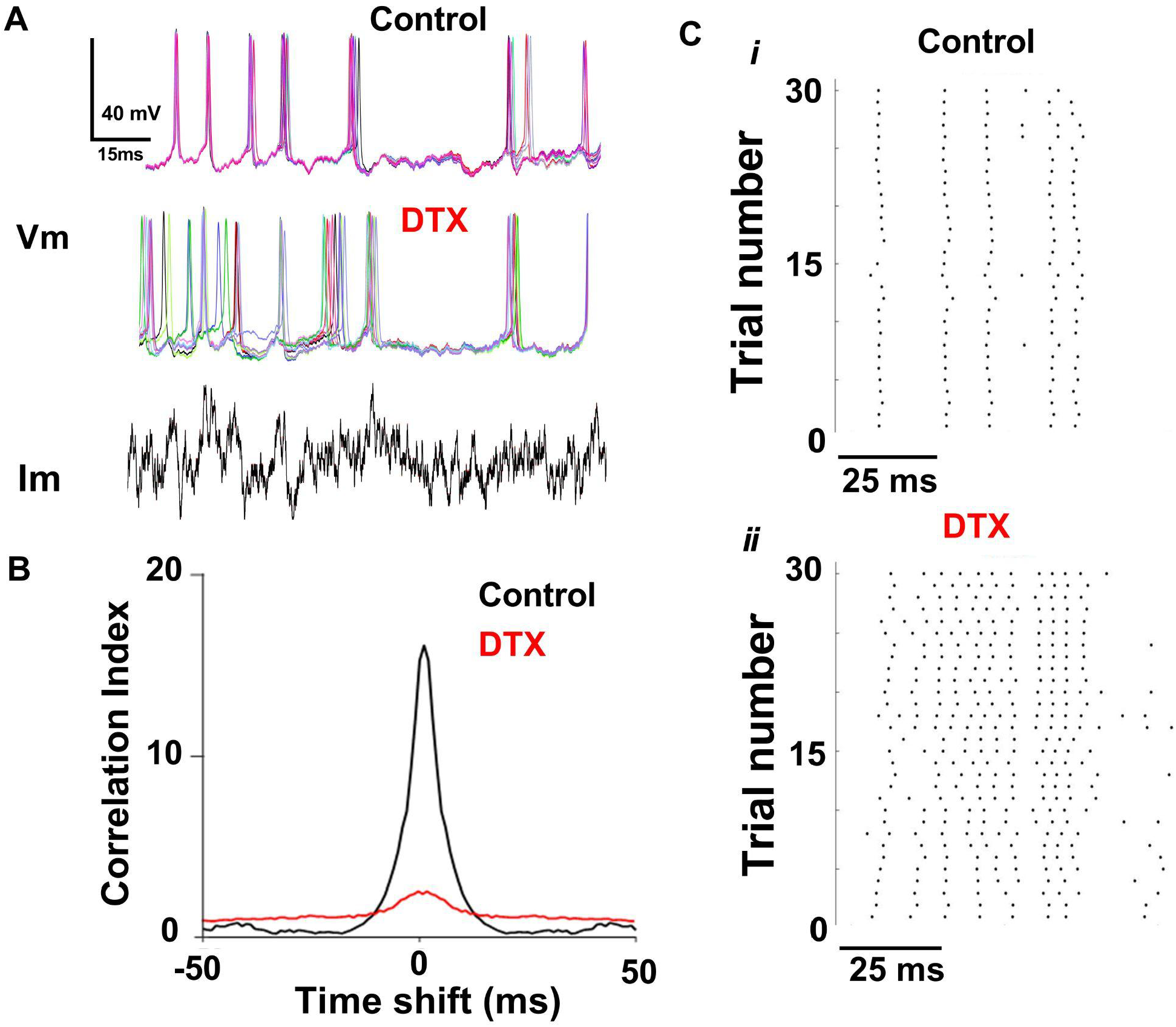
DTX reduced the spike timing reliability in a tonic II neuron in NA. A) Overlaid voltage traces (V_m_) for seven repetitions of an identical “noisy” current (I_m_) stimulus in control and 100 nM DTX conditions. Only the first 150 ms of the 2 second long stimulus is shown. B) Raster plots of all 30 trials for each condition, first 100 ms of response (onset at 0 ms). C) Spike timing reliability quantified with a shuffled autocorrelogram showed a reduced peak correlation index and broader distribution for responses in the presence of DTX.

**Fig. 4.**
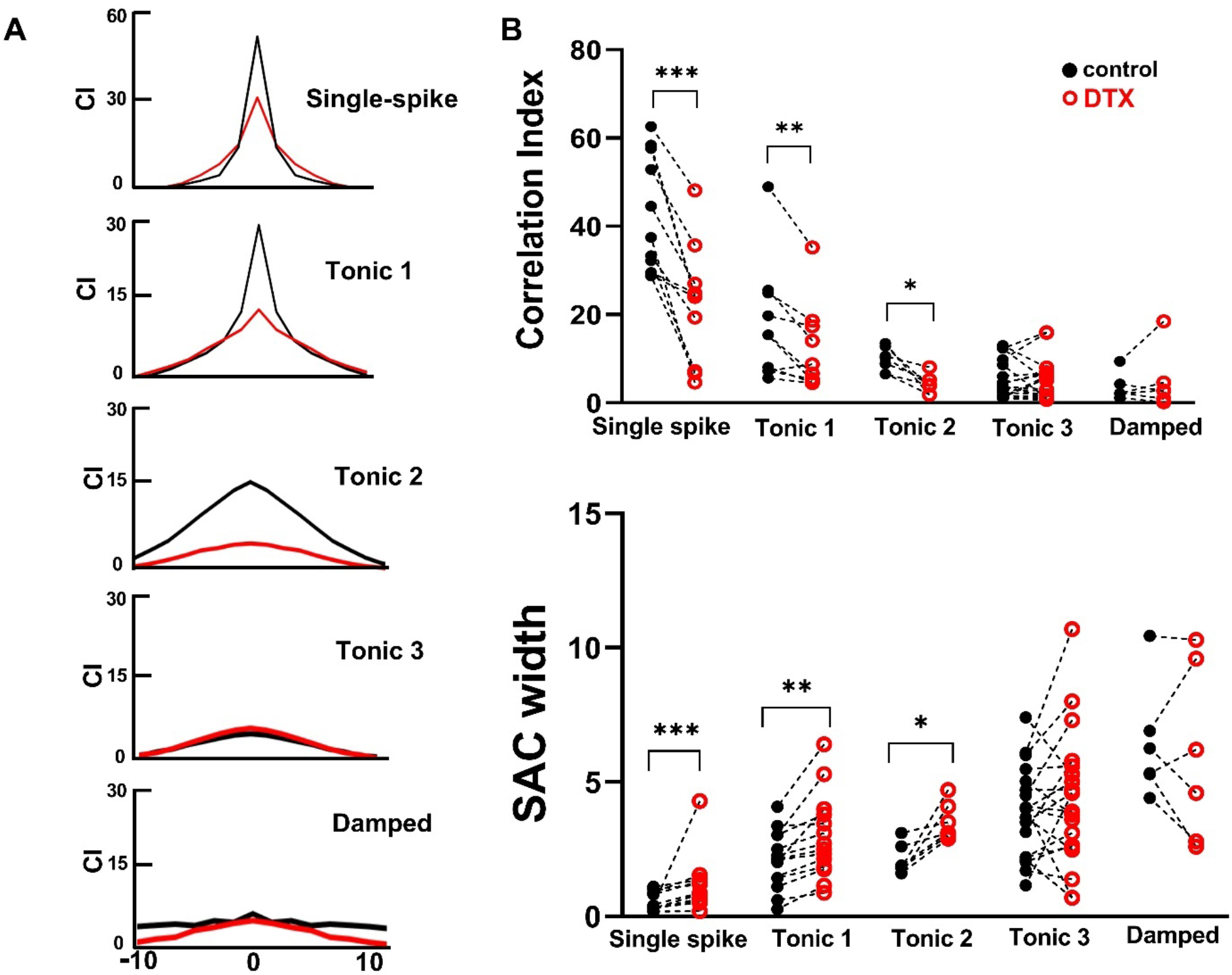
DTX reduced the spike timing reliability in cell type-specific manner. (A) Averaged SAC across all neurons recorded for each of the 5 cell types (control, black, DTX, red). (B) Summary data of SAC parameters peak correlation index (CI)(top) and full width at half-maximum height (FWHM) (bottom). Peak CI decreased in single-spike, tonic I, and tonic II neurons (P < 0.001, P = 0.0028, p = 0.0087, respectively, Wilcoxon’s test) while corresponding FWHM values increased (P < 0.001, P < 0.001, P = 0.028).

Another method to quantify the similarity (or dissimilarity) of the spiking responses before and after DTX application is to measure the mutual information (MI) contained in the two sets of spike trains (see Methods)(37, 38). MI quantifies the predictive value of one data set toward the other in bits, where 1 indicates complete certainty and zero indicates complete independence between the data sets. The MI between control and DTX trials was low in single-spike, tonic I, and tonic II neurons (Fig. 5, red markers and bars), indicating a high degree of dissimilarity. The same analysis applied to separate experiments that used sham drug application yielded an MI close to 1, significantly higher than for DTX experiments (P < 0.001, P = 0.007, P = 0.028 respectively, Wilcoxon’s test). For tonic III neurons, the mutual information between control and DTX spike trains was somewhat higher than for other neurons but was similar to that during sham experiments, suggesting DTX had no specific effect on these neurons (P = 0.708) (Fig. 5). Together, these spike train timing analyses suggest that Kv1 channels are crucial for the specific and reliable encoding of dynamic stimuli in a subset of NA neurons.

**Fig. 5.**
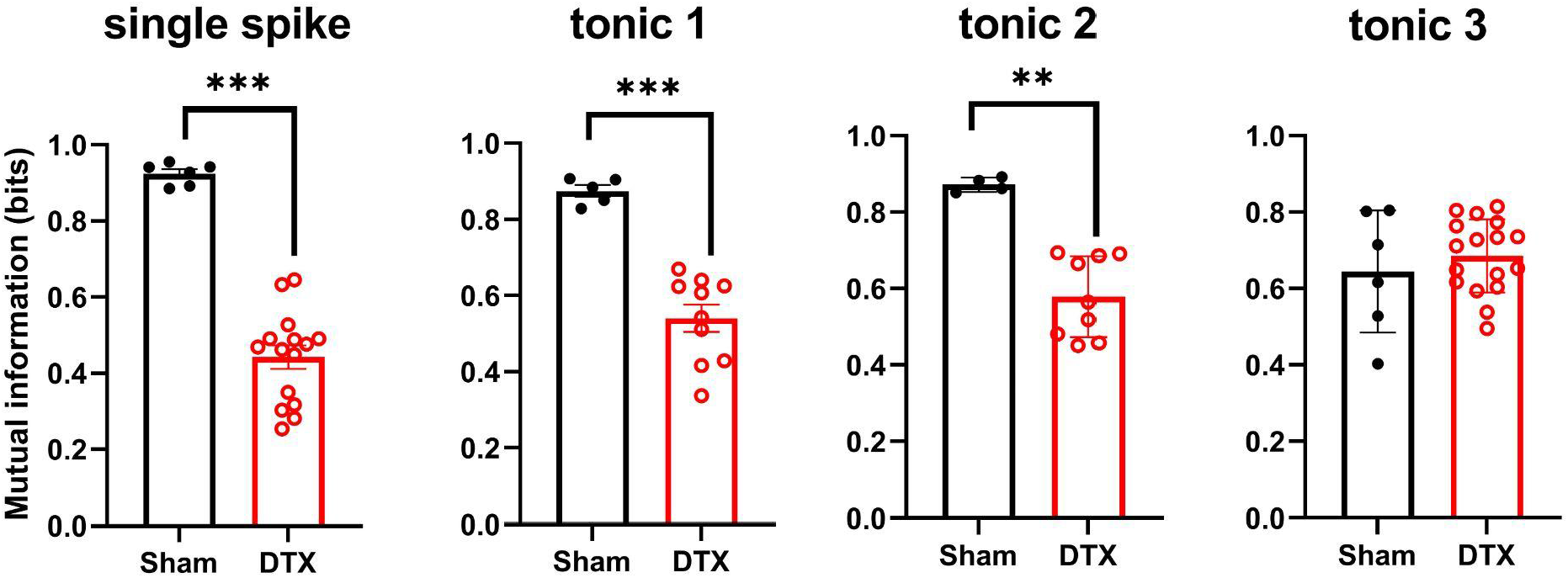
Mutual information (MI) between spike trains by cell type. MI between control spike trains and spike trains following sham solution change was high (black markers, bars) indicating high similarity. In contrast, MI between spike trains from control versus DTX trials was significantly lower for single-spike, tonic I, and tonic II neurons but not for tonic III neurons (Mann-Whittney test, P < 0.001, P < 0.001, P = 0.003 and P = 0.707, respectively).

### Kv1 channel expression drives fluctuation sensitivity by contributing to an adaptive spike threshold

A key feature of coincidence detector neurons is their selective responsivity to highly correlated synaptic inputs that cause a rapid rise in the postsynaptic voltage. To determine whether DTX affected the selectivity of NA neurons to rapid changes in their inputs, we measured the spike-triggered average current (STA) of the stimulus preceding each action potential. The peak-to-trough amplitude of the STAs were reduced by application of DTX in the single spike, tonic I, and tonic II neurons (single spiking control amplitude = 0.11 ± 02 nA, DTX amplitude = 0.044 ± 0.01 nA, n = 11, P < 0.001, Wilcoxon t-test; tonic I control amplitude = 0.032 ± 0.005 nA, DTX amplitude = 0.019 ± 0.001 nA, n = 9, P = 0.036; tonic II control amplitude= 0.031 ± 0.02 nA, DTX amplitude = 0.0185 ± 0.0015 nA, n = 8, P = 0.014) (Fig 6). In contrast, the peak-trough amplitudes of the STAs for the tonic III and damped neurons were unaltered (tonic III amplitude = 0.015 ± 0.0013 nA, DTX amplitude = 0.016 ± 0.003 nA, n = 16, P = 0.73 ; damped control amplitude = 0.013 ± 0.003 nA, DTX amplitude = 0.013 ± 0.003 nA, n = 5, P = 0.95). A large STA amplitude indicates that the neuron will only fire to rapidly rising inputs, so a reduction in STA amplitude implies a corresponding reduction in selectivity.

**Fig. 6.**
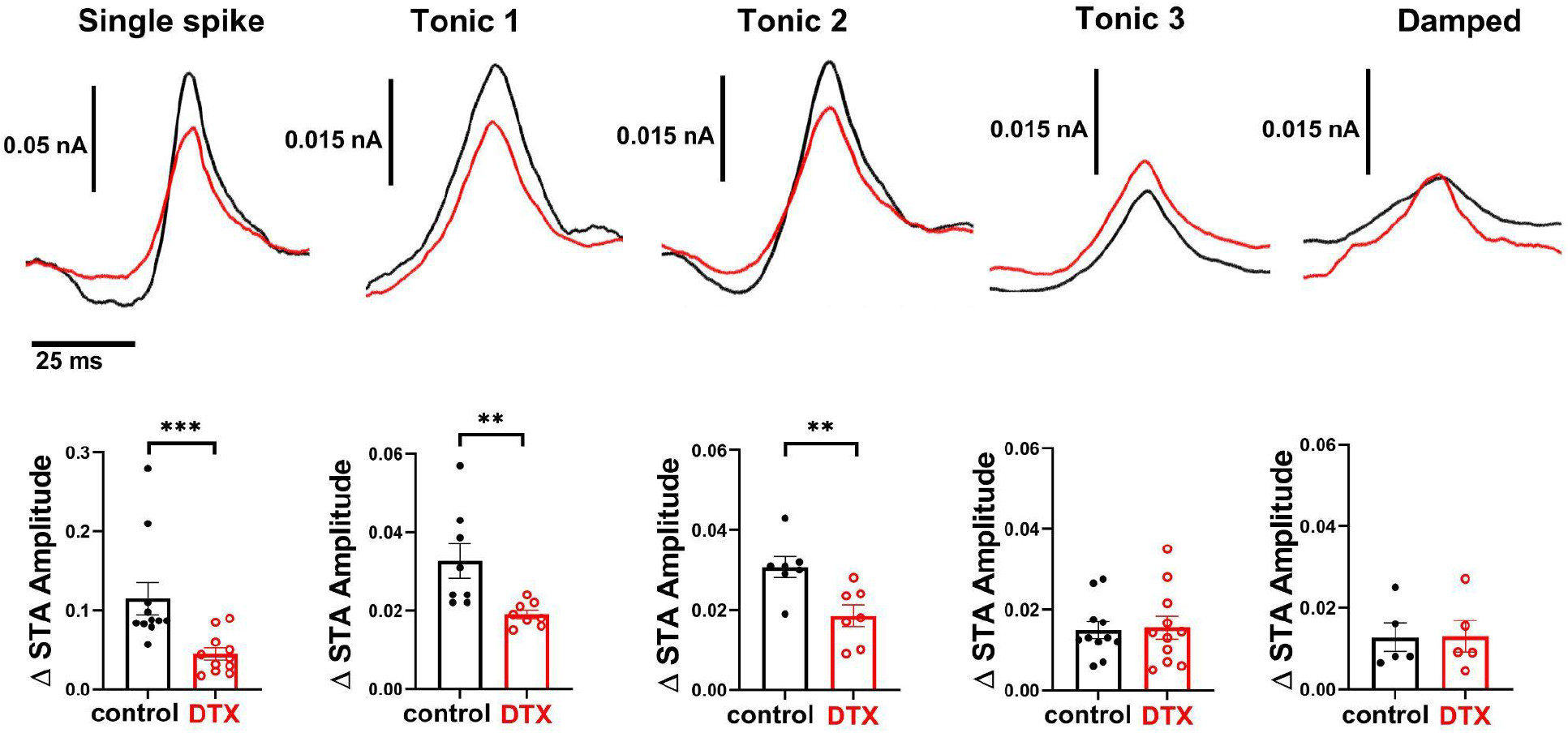
DTX reduced the high-pass response selectivity in a subset of NA neurons. Top row: spike-triggered averages (STA) for each neuron type before (black traces) and after (red traces) DTX application. Traces are grand averages of the population for each cell type. Bottom row: summary statistics of peak-to-trough amplitude of STA waveforms from individual neurons. STA amplitudes were significantly reduced in the presence of DTX for single-spiking, tonic I, and tonic II neurons (P < 0.001, P = 0.036, P = 0.014, respectively, Wilcoxon’s t-test). Tonic III and damped neuron STAs were unchanged.

Previous studies of the input-output function of the repetitively firing neurons in NA showed that sensitivity to rapidly rising inputs leads higher firing rates to stimuli containing larger fluctuations (19, 26, 28). Neurons that showed the greatest fluctuation sensitivity have been dubbed ‘differentiators’, in contrast to ‘integrator’ neurons whose firing rates reflected the mean stimulus level regardless of the size of the noise fluctuations around the mean (39). To determine whether G_KLT_ affected fluctuation sensitivity, we measured the *f*-I curves using current steps with two noise fluctuation amplitudes (2σ, 4σ and 8σ) before and after DTX application. In 15 tonic firing NA neurons classified as differentiators (one control example shown in Fig. 7Ai), DTX application elevated firing rates overall (the maximum measured firing rate increased from a mean across 15 neurons of 68.4 Hz in control to 83.7 Hz with DTX, data not shown). However, it also reduced the relative firing rate enhancement to larger current noise fluctuations compared to smaller noise fluctuations (Fig. 7Aii). These noise *f*-I curves were quantified by measuring the normalized difference in maximum firing rate (MaxFR Index) between the low noise (2σ) and high noise (8σ) and the normalized firing rate difference over the entire curve (Area Difference Index, ADI) (see Methods). Both these measures are larger in differentiators than in integrators (which are close to the origin, Fig. 8Ai) and decreased significantly with DTX (Fig. 7Bi-Biii MaxFR Index control = 0.388 ± 0.152, Differentiator DTX = 0.199 ± 0.146, n = 15, P < 0.001, Student’s t-test; ADI control = 0.455 ± 0.175, DTX = 0.273 ± 0.158, P < 0.001). In contrast, application of DTX had little effect on the noise *f*-I curves in 5 tonic firing NA neurons classified as integrators (Fig. 8A, MaxFR Index control = 0.035 ± 0.034, DTX = 0.029 ± 0.0581, n = 5, P = 0.42, Student’s t-test; ADI control = 0.052 ± 0.072, DTX=0.043 ± 0.083, P = 0.39). It should be noted that this reduction cannot be explained by a simple proportional increase in all firing rates, which would have had no effect on the normalized indices. Thus blocking G_KLT_ with DTX selectively reduced the fluctuation sensitivity by the differentiator neurons, causing them to behave more like an integrator neuron.

**Fig. 7.**
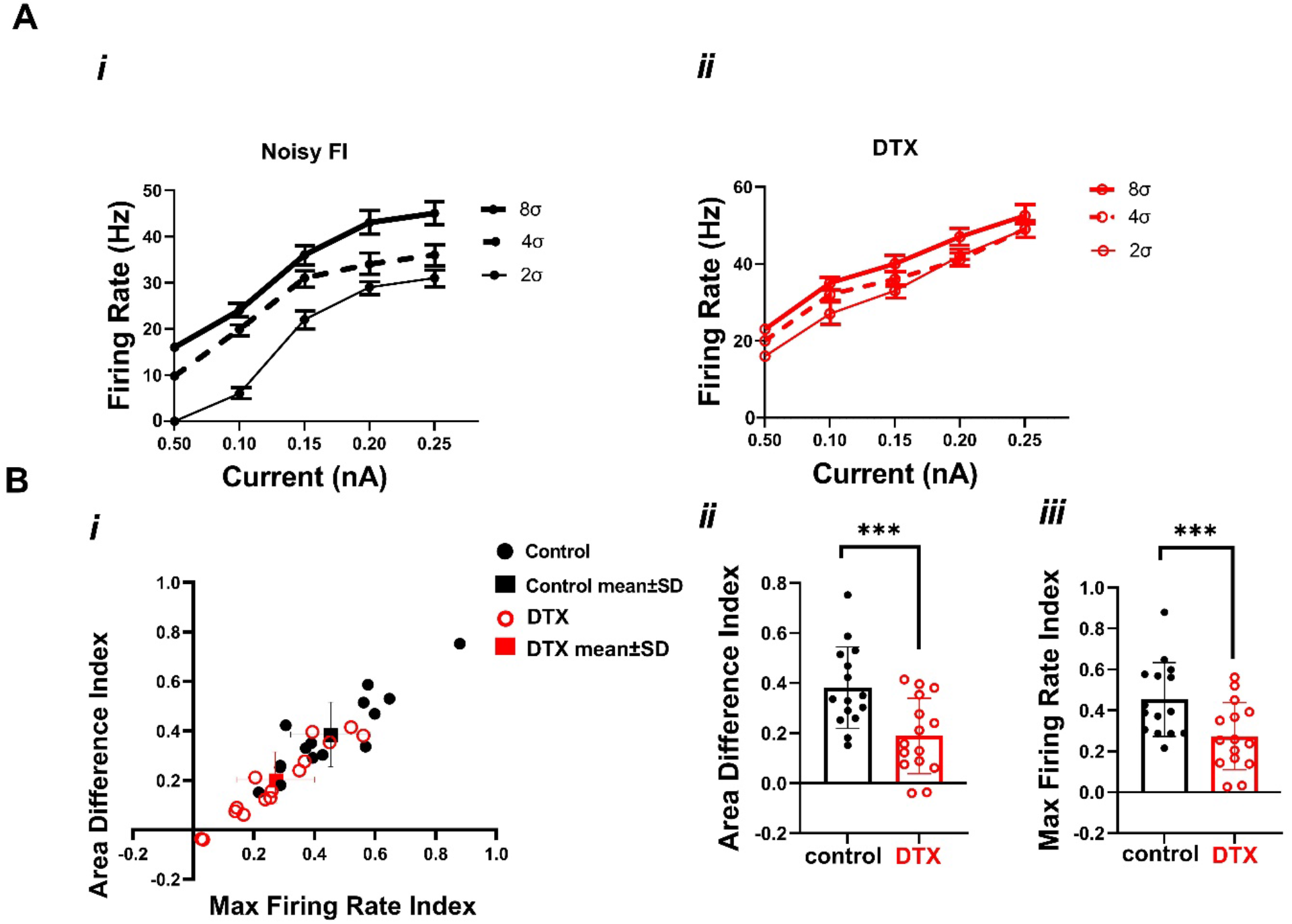
DTX diminished fluctuation sensitivity in differentiators. **(A)**An example of one differentiator neuron’s input-output function with 3 levels of noise before (*i*) and after DTX application (*ii*). (B) Summary plot of f-I curve metrics for 15 differentiator neurons (Bi, black solid and red open markers represent before and after pairs; square markers with error bars are group mean±SD). (B*ii*) Area Difference Index (control ADI = 0.391 ± 0.17, DTX ADI = 0.187 ± 0.15) and (B*iii*) maximum firing rate index (control ΔmaxFR = 0.422 ± 0.19, DTX ΔmaxFR= 0.261 ± 0.15) are reduced upon DTX application.

**Fig. 8.**
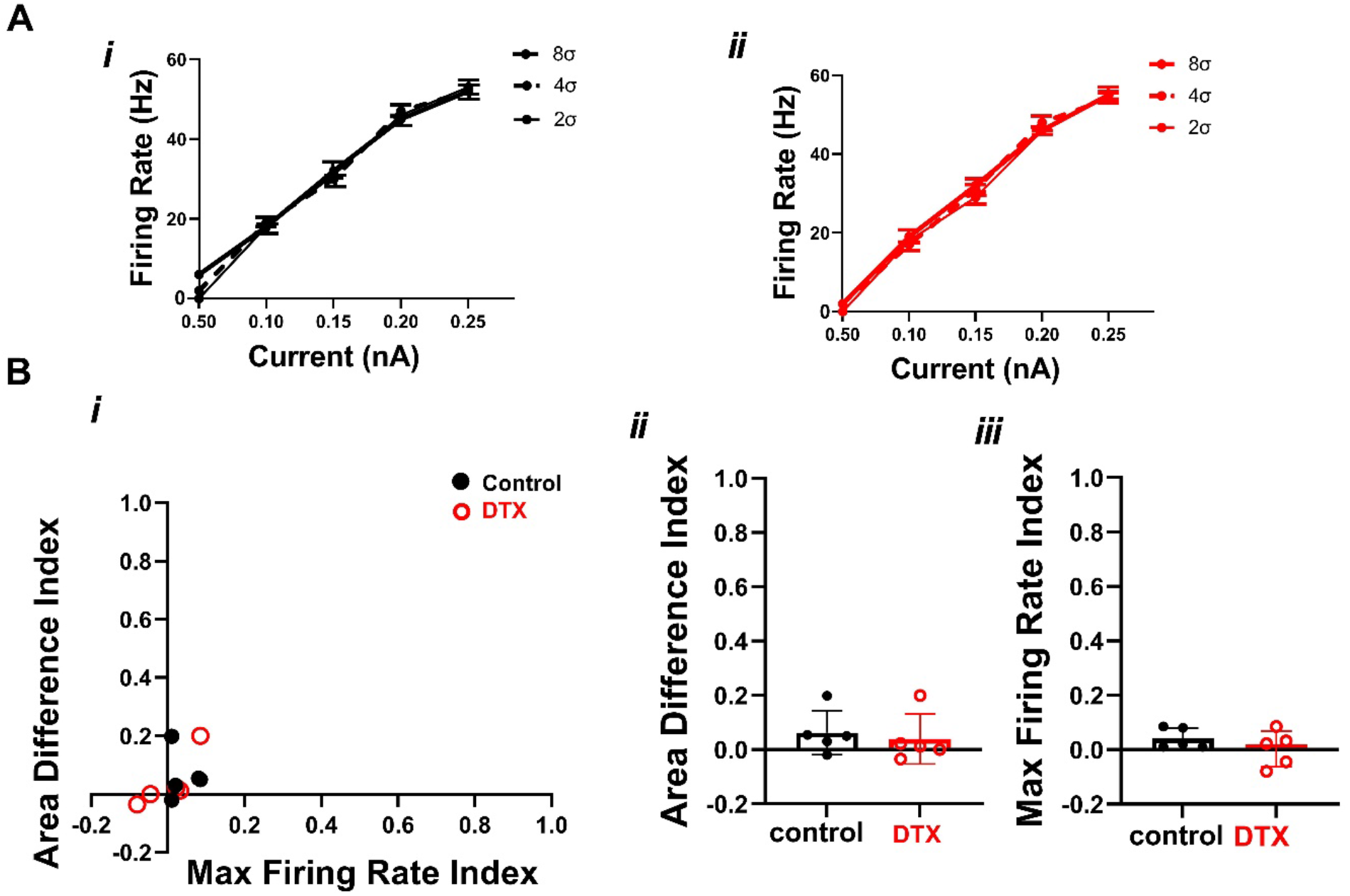
DTX had no impact on fluctuation sensitivity metrics in integrators. Panels as in Fig. 9. An example of one integrator neuron’s input-output function with 3 levels of noise before (Ai) and after DTX application (Aii). (B) Summary plot of f-I curve metrics for 5 integrators neurons indicated no DTX effects. (Bii) Area Difference Index (control ADI = 0.042 ± 0.14, DTX ADI = 0.032 ± 0.025) and (Biii) maximum firing rate index (control ΔmaxFR = 0.036 ± 0.017, DTX ΔmaxFR = 0.008 ± 0.072) show no changes with DTX.

The primary mechanism underlying the fluctuation sensitivity and high pass behavior has been proposed to be an adaptive spike threshold (31, 40, 41). Sodium channel inactivation is a key ion channel mechanism to achieve the adaptation (31), but other conductances that are active in the subthreshold regime have the potential to influence threshold (32, 34, 39, 42). During adaptation, the voltage threshold dynamically varies depending on the mean voltage, the rate of rise in the voltage, and the time since a preceding action potential. We asked whether the effects described above are consistent with a role for G_KLT_ in spike threshold adaptation. To measure spike threshold adaptation, we measured threshold variation during noisy current drive, as previously described in NA neurons (28). With DTX application, the variation in the threshold to the large noise stimulus (8*σ*) was reduced (Fig9A), and the spike waveforms were more stereotyped (Fig9B, C). Across 15 NA tonic firing neurons, DTX significantly reduced the threshold variation across a range of mean current amplitudes (2-way ANOVA, main effect by drug, [F(20, 345)= 10.95, P =0.001]; P = 0.003 at 0.20 nA, P=0.0021 at 0.25 nA, P<0.001 at 0.30 and 0.35 nA, Sidak’s multiple comparisons test)(Fig. 9D). These data show that consistent with the reduction in filtering, DTX affects fluctuation sensitivity at least in part due to reduced threshold adaptation. These results suggest that Kv1 channels are involved in threshold adaptation at higher noise levels, possibly in conjunction with sodium channel inactivation.

**Figure 9.**
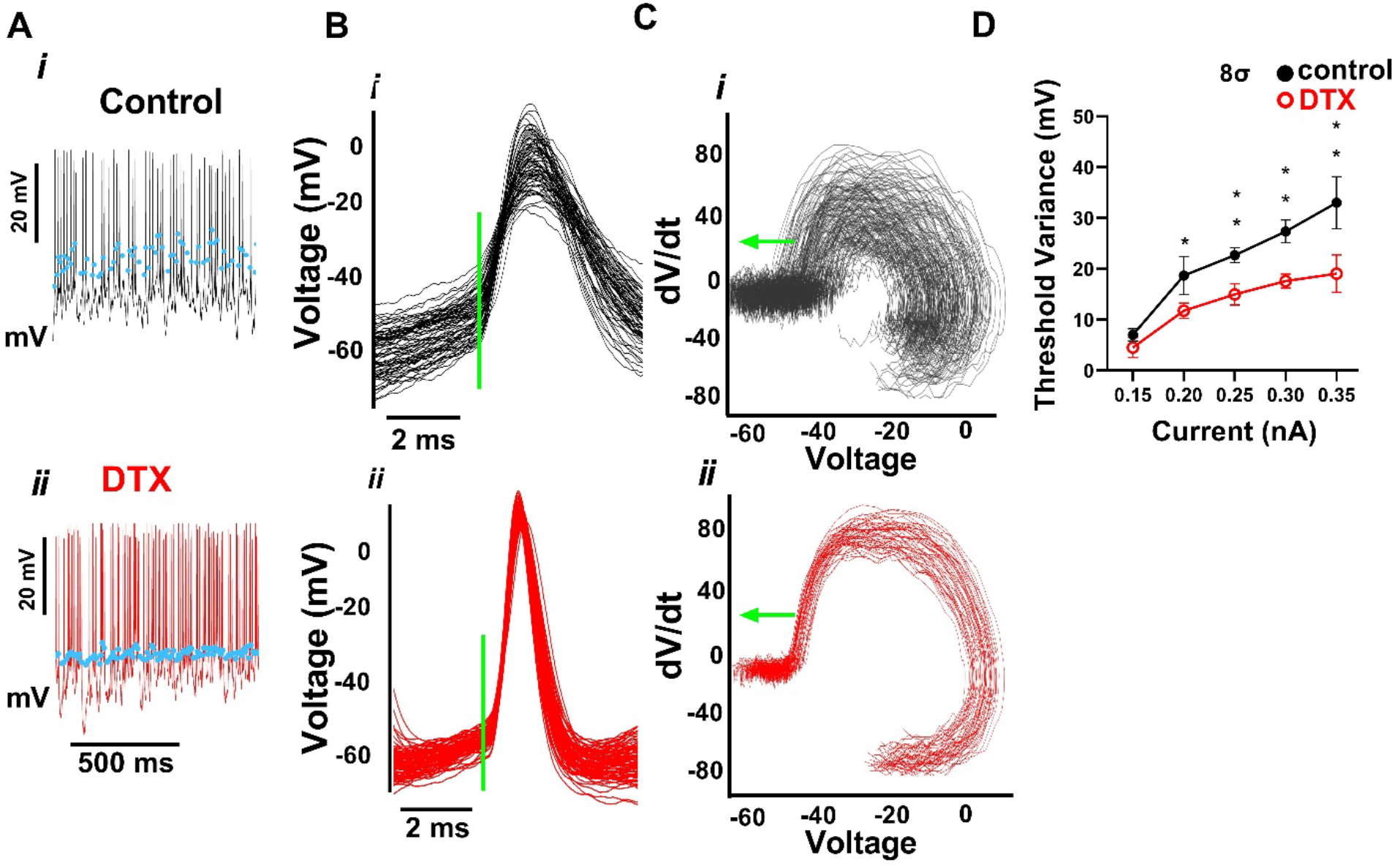
DTX altered the dynamics of action potential initiation and threshold variability during periods of high current fluctuation. (A) Example spike trains with spike thresholds indicated by blue dots in a control (*i*) and post-DTX application (*ii*). (B) Action potentials overlaid for control (*i*) and DTX (*ii*) trials aligned on spike threshold (green bar). (C) Voltage derivative versus voltage for APs in panel B for control (*i*) and DTX (*ii*). dV/dt threshold is labeled with a green arrow. (D) Threshold variability was significantly diminished in the 8*σ* noise fluctuation level from 0.2 nA and greater (2-way ANOVA, main effect by drug, [F(20, 345)= 10.95, P =0.001]; 0.2 nA: P = 0.0085; 0.25 nA: P =0.0041; 0.3 nA: P=0.001; 0.35 nA: P =0.001; Sidak’s multiple comparisons test).

## Discussion

The role of the low-threshold gated potassium conductance (G_KLT_) in the temporal processing of auditory signals has been extensively studied. The G_KLT_ conductance is attributed to the expression of members of the Kv1 family of voltage gated potassium channels, particularly Kv1.1 and Kv1.2, that are prevalent throughout the auditory brainstem structures of birds and mammals (14, 15, 22, 43). Distinct from the Kv3 family associated with a high threshold conductance responsible for action potential repolarization (44–46), in vitro studies showed that GKLT activates at membrane potentials near rest and is responsible for the single spiking behavior and outward rectification characteristic of many auditory neurons (5, 6, 47–49), Activation of G_KLT_ results in a short membrane time constant that prevents the temporal summation of synaptic potentials (9, 50, 51). Thus the G_KLT_ conductance is a key component to phase locking to the fine temporal structure in the acoustic signal and high fidelity sensory encoding in the cochlear nucleus NM (3, 23, 52). In the avian interaural time difference (ITD) circuit, dynamic activation of this conductance is crucial for the spike initiation properties that underlie the coincidence detection in the NL (5, 6, 8, 22, 32, 39, 53–56). Similar effects can be observed in the analogous mammalian circuit (42, 57–61).

Pharmacological and immunohistochemical studies, however, have suggested that Kv1 channels are not exclusively expressed in these timing pathways but are also present in the sister cochlear nucleus, NA (10, 22, 30). In this study, we found that pharmacological blockade of G_KLT_, with the Kv1.1/1.2 specific antagonist DTX, dramatically altered the firing phenotypes of some NA neuronal subtypes but not others. Specifically, the single-spiking neurons and the tonic I and tonic II phenotypes lost their unique onset or bursting firing patterns, while the tonic III phenotype that only displayed regular spiking was unchanged. Additionally, we found that the temporal reliability of firing, high-pass filtering characteristics, and spike threshold adaptation were all partially reliant on the DTX-sensitive conductance. These results suggest that Kv1 channels make important contributions to the intrinsic physiological diversity and functional heterogeneity of NA neurons.

### Ionic basis of electrophysiological diversity in NA

Previous work showed that DTX application to single spiking NA neurons converted them into repetitive firing neurons (22). More surprising was the finding that DTX also caused an increase in firing rate in NA tonic firing neurons. We confirmed and extended these results to show that DTX-sensitive changes in firing pattern and rate are related to the physiological classifications in the chick embryonic brain stem described previously (25). The DTX effects were limited to tonic firing neurons that show bursting behavior at lower current stimulus levels. We conclude that the burst termination (tonic I) and delayed burst (tonic II) behaviors both require the presence of a DTX-sensitive conductance, but it remains unclear what causes the differences in their response patterns. One explanation could be a difference in the activation or inactivation kinetics of the Kv1 channel conductance; a slow inactivation of I_KLT_ has been shown in NM neurons (62). Since two isoforms, Kv1.1 and Kv1.2, are both DTX-sensitive, there may be differences in subunit compositions or modulation. Alternatively, there may be some other subthreshold conductance present in one type of tonic neuron that interacts with the DTX-sensitive current (e.g., an A-type current that could enhance early bursting (33, 34). Two neuron types in NA appeared unaffected by DTX: tonic III neurons which best represent canonical rate-coding integrators, and damped neurons, a potential immature phenotype (22, 29). In the presence of DTX, the blockade of bursting resulted in responses by all three tonic firing subtypes more closely resembling each other. Because these subtype differences have not been observed in the hatchling (22), it is possible that the increase in excitability and reduction in phenotypic diversity that occurs with development could be due to a downregulation of G_KLT_. Examining these neuronal subtypes is important because computational approaches have shown that increased population heterogeneity is directly correlated with increased information encoding capacity (27).

### Neurons in the NA utilize Kv1 channels to encode the temporal dynamics of their inputs

Multiple features demonstrated as archetypal in NA appear to be at odds with a presumed role for G_KLT_ in temporal processing: multi-spiking intrinsic firing properties (1, 22, 26), poorer phase-locking in vivo to the acoustic fine structure, and a role in interaural level coding (3, 16, 23). However, a more complex picture of sound processing in NA has been emerging. First, the heterogeneity of in vivo response properties and morphology suggested that NA is not a monolithic rate-coding nucleus, but a diverse population of neurons (21, 25), some of which show temporal sensitivity, dynamic feature selectivity, and auditory envelope processing (19, 20, 26, 28). Second, in vitro studies showed a diversity of intrinsic features across the population, some of which also demonstrated greater temporal sensitivity than others (25–27). In another study, the differentiator-type intrinsic properties of a subset of NA neurons could account for the intensity-dependent bandpass behavior in vivo (19). In their model, the high-pass filtering due to active intrinsic membrane properties of the tonic-firing, coincidence-detector-like neurons combined with the low-pass filter behavior of the synaptic inputs and passive membrane properties resulted in the overall bandpass filtering. Interestingly, variation in the properties across the neuronal population produced an array of bandpass properties spanning the physiological response space. The results presented in the present study provides direct evidence that variations in Kv1 conductances contribute to the variations in temporal processing properties in vitro; it remains to be tested whether a mature circuit in vivo would also require these properties to encode spectrotemporal auditory information. Neurons with high reliability should be capable of encoding more rapidly changing components of the acoustic signal, such the onsets and gaps that are key features of speech and other communication signals. Neurons with less reliability are likely to be important for encoding the more slowly changing aspects of the acoustic signal, such as slower envelope variations or ongoing interaural level cues. Together, these studies suggest that NA neurons function on a spectrum of operating modes, ranging from pure integrators to pure coincidence-detectors (34).

### Spike threshold adaptation in NA neurons may rely on the low threshold potassium conductance

The operating mode diversity observed across NA specifically appears to arise from a mechanism known as spike-threshold adaptation, a phenomenon by which the voltage threshold is modulated by spike history and membrane voltage mean and rate of rise (19, 63–65). This form of adaptation is the probable mechanism underlying the selective responsiveness of NA neurons to rapid rises in the postsynaptic voltage, acting like a high-pass filter and enhancing the firing response to temporally modulated inputs. Spike threshold adaptation is a process that occurs throughout the brain, particularly in cortical pyramidal neurons, and auditory brainstem neurons (19, 28, 31, 32, 66, 67). In NA, threshold adaptation drives NA neurons to encode amplitude modulations with higher fidelity than auditory nerve fibers (19) as well as diversifies their operating modes (28). Phenomenological models based on sodium channel inactivation as a mechanism were successful at recapitulating the firing responses (19, 28), however these models use abstract conductance representations that do not specifically rule out other non-sodium, subthreshold conductances, such as the Kv1 channel. In a study of cortical neurons, blockade of Kv1 channels with DTX largely eliminated spike threshold adaptation (32). Our results suggest a similar role for Kv1 in NA neurons.

### Functional analogs of NA neurons in the mammalian cochlear nucleus

The mammalian auditory brain stem is not anatomically homologous to the avian auditory brain stem. The mammalian ventral cochlear nucleus (VCN) has functionally similar cell types to those in NA, but there does not appear to be a one-to-one match for each type. For example, the spherical bushy cells of the VCN are analogous to the avian NM neurons in that both have high levels of Kv1 expression, receive calyceal inputs from auditory nerve fibers, generate precise spiking phase-locked to the fine structure in auditory signals and project to analogous coincidence detection circuits (68). The T-stellate cells of the VCN appear to be the closest functional analog to NA neurons, in that they have reduced phase-locking relative to auditory nerve inputs, repetitive intrinsic firing properties, encode envelope and intensity information for spectrotemporal coding, and project directly to the midbrain inferior colliculus (1, 12, 69–74). Identified T-stellate neurons lack any significant low threshold potassium channel conductance and are insensitive to DTX (75–78). These results suggest that the T-stellate neurons of the VCN are not functionally analogous to the DTX-sensitive neurons in NA (tonic I/tonic II subtypes), but instead may be more similar to the DTX-insensitive, integrator like neurons of NA (tonic III subtype). Another study, using acutely dissociated neurons and voltage clamp, suggested there was a population of neurons in the VCN with intermediate levels of DTX-sensitive, lower-threshold conductances, possibly corresponding to the radiate (or D-stellate) type of multipolar neurons, a less common type of repetitively firing neuron in the AVCN that projects to the dorsal cochlear nucleus (72, 79–81). The recent discovery in the VCN of a third stellate neuron population, dubbed L-stellate, which appears to provide narrowband inhibition and operates within a feedback loop with T-stellate neurons (82) further suggests there may be greater phenotypical diversity than previously appreciated. How these cell types may correspond to NA cell types remains to be seen. These multi-spiking neurons in VCN can entrain to high frequency stimuli in vitro, but the reliability of responses to the noisy, temporally modulated stimuli we describe in the present study have only been tested in DCN neurons (36). Comparisons between NA and VCN are complicated by the fact that VCN contains a dense network of lateral connectivity and local inhibition, while NA appears to be a strictly feedforward circuit without local inhibition, so their feature selectivity and computational roles are likely to be quite dissimilar. Also limiting comparisons of intrinsic properties across studies are species differences (chick vs guinea pig vs mouse) and the possible confounding effect of temperature on the kinetics of the ion channels (78, 83).

### Summary

In summary, we have demonstrated that dendrotoxin-sensitive conductances are prevalent in a subpopulation of NA neurons and are critical for the temporal response properties in these neurons. By enhancing spike threshold adaptation, this conductance underlies the spike timing reliability, high-pass membrane filtering, and fluctuation sensitivity properties of the differentiator subtypes. The regulation of the Kv1 family of channels may be a key driver of intrinsic electrophysiological diversity in the avian cochlear nucleus and contribute to spectrotemporal auditory feature selectivity in vivo.

## Acknowledgements

Support for this research was provided by NIH grant R01DC10000. The authors thank Felix Bartsch for his helpful advice and MATLAB code for the mutual information analysis.

